# Effects of tendon viscoelasticity in the distribution of forces over sutures in a model tendon-to-bone repair

**DOI:** 10.1101/2021.11.09.467963

**Authors:** Yuxuan Huang, Ethan D. Hoppe, Iden Kurtaliaj, Victor Birman, Stavros Thomopoulos, Guy M. Genin

**Affiliations:** NSF Science and Technology Center for Engineering Mechanobiology, Department of Mechanical Engineering and Materials Science, Washington University, St. Louis, MO, USA; Department of Mechanical and Aerospace Engineering, Missouri University of Science & Technology, Rolla, Missouri, USA; Department of Orthopedic Surgery, Columbia University, New York, NY, USA

## Abstract

Tears to the rotator cuff often require surgical repair. These repairs often culminate in re-tearing when sutures break through the tendon in the weeks following repair. Although numerous studies have been performed to identify suturing strategies that reduce this risk by balancing forces across sutures, none have accounted for how the viscoelastic nature of tendon influences load sharing. With the aim of providing insight into this problem, we studied how tendon viscoelasticity, tendon stiffness, and suture anchor spacing affect this balancing of forces across sutures. Results from a model of a three-row sutured re-attachment demonstrated that optimized distributions of suture stiffnesses and of the spacing of suture anchors can balance the forces across sutures to within a few percent, even when accounting for tendon viscoelasticity. Non-optimized distributions resulted in concentrated force, typically in the outermost sutures. Results underscore the importance of accounting for viscoelastic effects in the design of tendon to bone repairs.

## 1 Introduction

Rotator cuff tears are widespread sources of pain and disability, especially amongst the elderly [1]. Massive tears usually require surgical repair to assist healing. However, the surgical repair often does not lead to tissue regeneration and has poor outcomes, with retear rates ranging from 11 to 94%[2–9]. Factors that can influence the post-surgery outcomes include patient age, the severity of the rotator cuff tear, and the rehabilitation protocol[10]. Improving rotator cuff surgical outcomes is increasingly important as the population ages and the demand for these repairs increases, and motivates efforts to develop new approaches for rotator cuff repair.

Standard rotator cuff repair techniques utilize multiple bone anchors that enable the surgeon to cinch the tendon to humeral head. Failure often occurs when the force on a suture reaches the value needed to pull the suture through tendon [11]. Our long-term goal is to improve suture repair techniques by varying individual suture stiffnesses and locations of sutures to balance these forces and evenly distribute them across individual sutures in the tendon. Although a large literature exists on this problem [4, 12–15], very few analytical approaches have been developed [16, 17], and none of which we are aware account for tendon viscoelasticity. We therefore developed a model to explore how tendon viscoelasticity may affect the optimization of repair.

Consideration of tendon viscoelasticity introduces time as a variable in the equations of motion. The amount of force necessary to produce a prescribed deformation increases if the loading interval is of short duration. Different viscoelastic models can be employed to describe the behavior of the tendon[18–21]. A necessary viscoelastic modeling is one of model accuracy versus simplicity [18]. Because our focus in this study was the development an understanding whether viscoelasticity might affect choices of suture patterns and designs, we opted to apply relatively simple models of tendon viscoelasticity in conjunction with the model of the rotator cuff repair. The model, which contained three rows of linear elastic sutures in a linear viscoelastic, one-dimensional tendon, was studied in the context of a broad range of loading and stiffness parameters. By studying the model in this parameter space, conditions emerged in which viscoelastic effects may become critical to repair design.

## 2 Methods

### 2.1 Model

A discrete viscoelastic shear-lag type model was studied for the purpose of illustrating how viscoelasticity affects the attachment in the context of tendon-to-bone attachment repair. The model consisted of three rows of idealized sutures affixed to immovable bone anchors at one end and to prescribed positions on a tendon at the other end (Fig. 1). The tendon was loaded with a horizontal force *F* (*t*) at one end, and was free at the other end. The magnitude of the force was ramped linearly from 0 to the peak value *F*_0_ at a constant rate 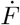, then maintained at a constant level *F*_0_ as the displacement *δ*_0_ of the loading point increased with time.

**Figure 1:**
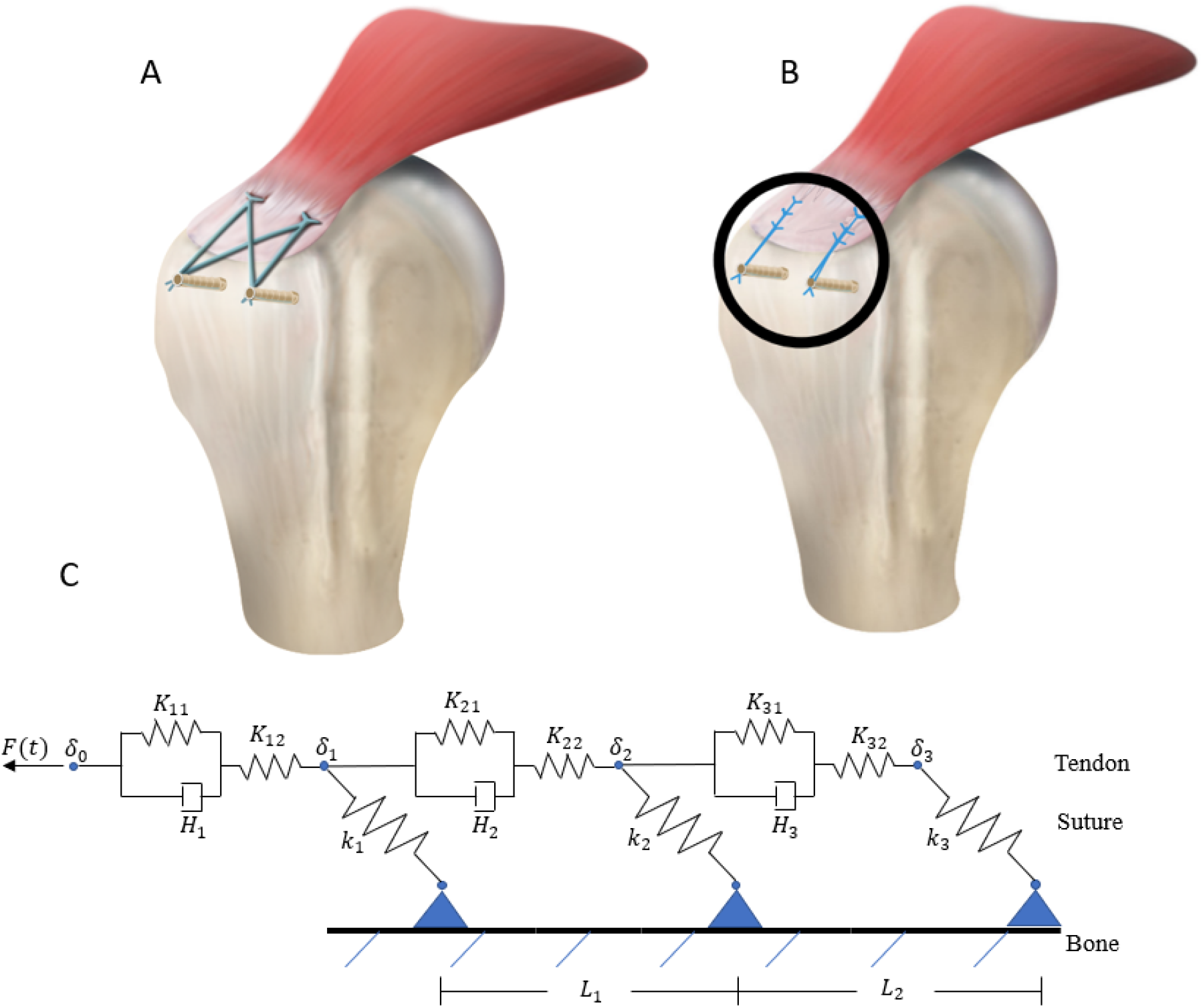
Tendon to bone repair model. **(A)** Traditional suture repair technique, with two rows of a single suture anchors in the tendon. Note that bone anchor screws, not shown, lie beneath the two suture insertion points in the tendon. **(B)** A three-row suture repair scheme, and the simplified model system analyzed (bone anchors are not shown). **(C)** The simplified system using 2D schematic. The bone-suture anchors are fixed at bone.

### 2.2 Constitutive Laws

Each row of sutures was treated as a linear spring with the stiffness of the *n*^*th*^ row from the end of the tendon denoted *k*_*n*_. Tendon segments were modeled as linear viscoelastic elements whose behavior is described by a three-parameter model (Fig. 2). This choice was made because a three-parameter model was the simplest that could replicate experimental observations in a meaningful way: the three-parameter model captures both creep and stress relaxation so that that models a material with some long-term, slow loading elasticity [22].

**Figure 2:**
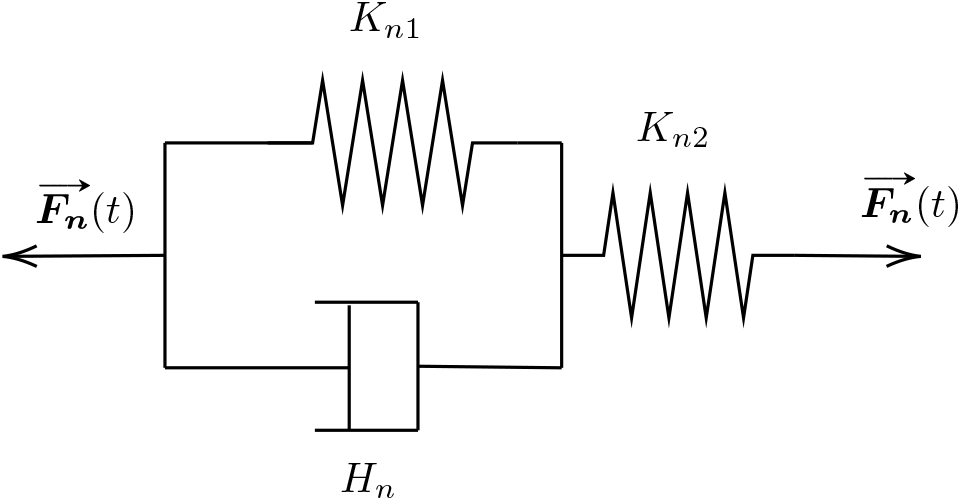
Three-parameter solid model used to represent individual segments of tendon. Each segment *n* had individual stiffness parameters *K*_*n*1_ and *K*_*n*2_, and viscosity parameter *H*_*n*_, which varied with segment length and spacing between rows of sutures. *F*_*n*_(*t*) is the force applied to the segment.

In Fig. 2, the system represents the *n*^*th*^ segment of tendon; segment 1 stretches from the loading point to the first suture row. *K*_*n*1_ and *K*_*n*2_ are the two stiffness parameters and the sum *K*_*n*1_ + *K*_*n*2_ was used to normalize suture row stiffness. In simulations in which spacings between suture rows were varied, the stiffnesses *K*_*n*1_ and *K*_*n*2_ both inversely proportionally to the anchor suture spacing. *H*_*n*_ is the viscosity of the tendon segment, which controls how fast the tendon reacts to the force.

To fit the baseline model to stress relaxation data for a rat supraspinatus tendon (dashed line, Fig. 3), a two timescale method was used, with separate fits for data in the 0-10 s and 10-550 s ranges. Short term parameters were *K*_*n*1_ = 11.5 N/m,*K*_*n*2_ = 8.51 N/m and *H*_*n*_ = 26.1N· s/m, and long term parameters were *K*_*n*1_ = 12.8 N/m,*K*_*n*2_ = 5 N/m and *H*_*n*_ = 1270 N· s/m [23].

**Figure 3:**
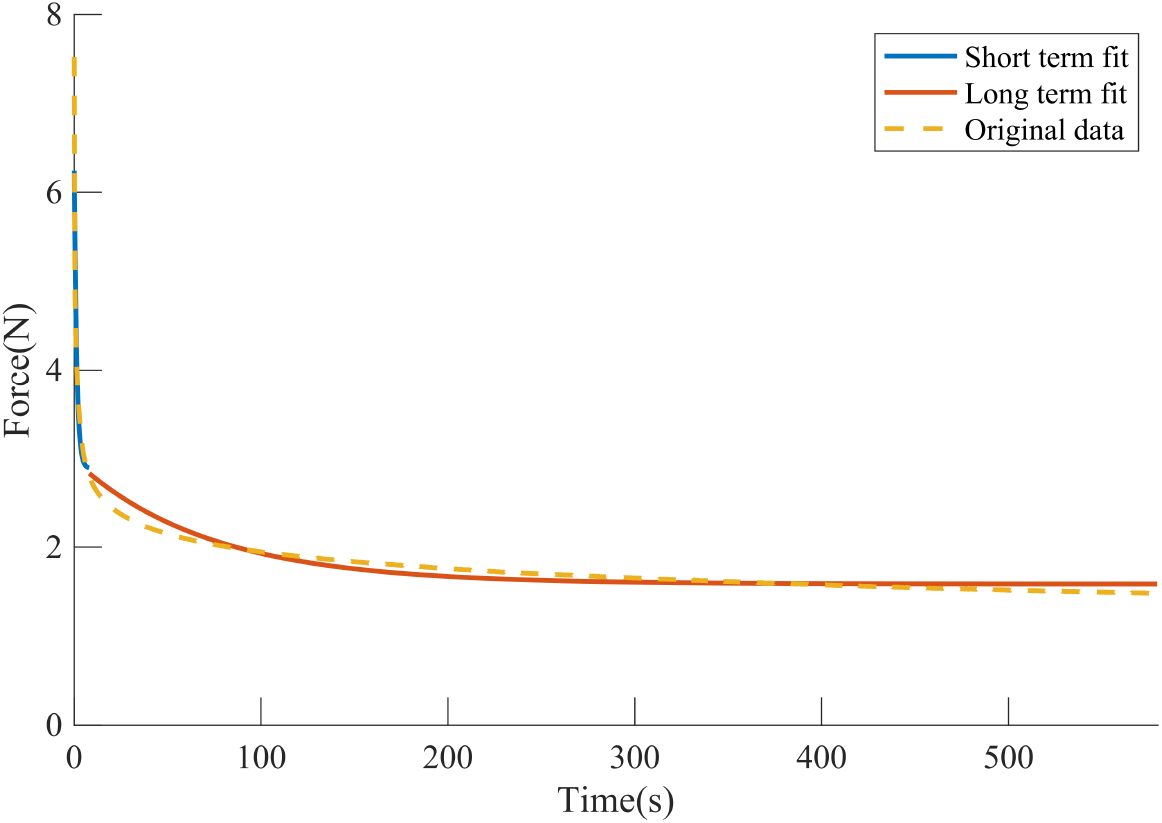
Curve fit of stress relaxation data of rat supraspinatus tendon using three-parameter model: short term and long term fit using the three-parameter model. Short term parameters: *K*_*n*1_ = 11.4 N/m,*K*_*n*2_ = 8.51 N/m and *H*_*n*_ = 26.0 N·s/m and long term parameters: *K*_*n*1_ = 12.8 N/m,*K*_*n*2_ = 5 N/m and *H*_*n*_ = 1270 N· s/m.[23]

### 2.3 Governing Equations

#### Kinematics

In the subsequent analysis, Δ_*n*_(*t*) is the deformation of segment *n* relative to its unloaded and undeformed reference length at time *t*, and was obtained as the difference between the displacements of two consecutive rows of sutures, Δ_*n*_(*t*) = *δ*_*n*_(*t*) − *δ*_*n*+1_(*t*) (Fig. 1). The rate of deformation of that segment was denoted by 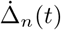.

#### Constitutive Equations

The three-parameter model force-displacement relationships for each tendon segment are derived below. Neglecting inertial terms, the force *F*_*n*_(*t*) in tendon segment *n* at time *t* in the parallel arrangement of the dashpot and spring can be written:

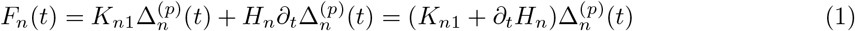

where *F* (*t*) is the force applied to the spring-dashpot system (Fig. 2), *H*_*n*_ is the dashpot viscosity, *K*_*n*1_is the parallel spring stiffness, 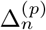 is the elongation corresponding to the parallel portion of the spring-dashpot system (*K*_*n*1_ and *H*_*n*_), and *∂*_*t*_() = *d*()*/dt*. This force equals to that on the spring of stiffness *K*_*n*2_ in the right side of the model (Fig. 3), so that 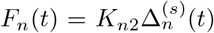, where 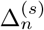 is the elongation of the spring *K*_*n*2_.

#### Equations of Motion

To write this in terms of the overall displacement of the tendon segment, 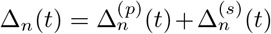, we note that 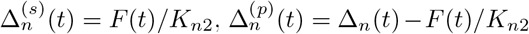, and write:

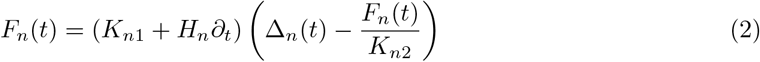

By reorganizing the terms one obtains:

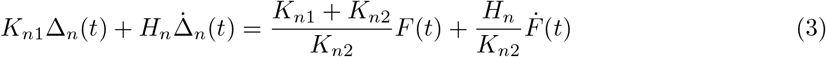

The force transmitted through the tendon to the *n*^*th*^ row of sutures is *k*_*n*_*δ*_*n*_ where *k*_*n*_ is the stiffness of the *n*^*th*^ row of suture. This is resisted by tension from the segment of tendon to the left, *F*_*n−*1_, and tension from the segment of tendon to the right, *F*_*n*_, so that so that *F*_*n*_ = *F*_*n−*1_ − *k*_*n*_*δ*_*n*_. This force *F*_*n*_ can be substituted into Eq. (3) for each suture row. The elongation Δ_*n*_ in Eq. (3) can be expressed as *δ*_*n*_ − *δ*_*n*+1_ for the corresponding tendon segment.

Additionally, the following condition of equilibrium must be satisfied at each time *t*:

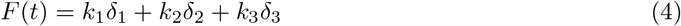

By utilizing the relationship above and applying Eqs.(3) and (4) to each tendon segment, we obtained three equations of motion that can be written in state space form as:

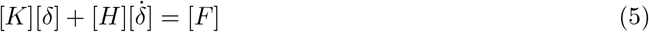

where [*δ*] and 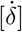 are vectors of nodal displacements and displacement rates, the stiffness matrix [*K*] is

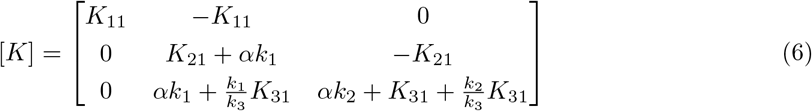

The damping matrix [*H*] can be written as

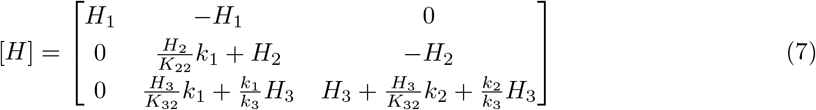

and the vector of nodal forces [*F*] is:

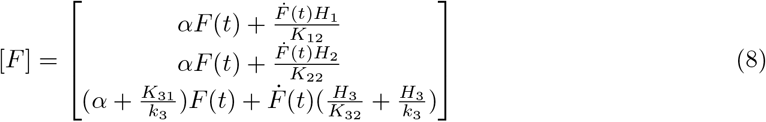

In the above equations, *α* = (*K*_*n*1_ + *K*_*n*2_)/(*K*_*n*2_); because the numerator and denominator vary inversely with tendon length, *α* is independent of the length of segments. Parameters *H*_*n*_, *K*_*n*1_, and *K*_*n*2_ depend on the lengths of the tendon segments. However, they share the following relationship with *L*_1_*/L*_2_:

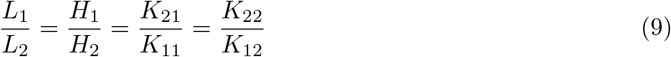

### 2.4 Numerical Methods

The form of the third row in equations (5)-(8) differed from that of the other two with more terms involved. The reason for this was that including this row based upon equilibrium of the third node caused the matrix [*K*] to become non-invertible, which resulted in a non-solvable system of equations. To circumvent this problem, the equation of the last row of sutures was replaced in the state space equations with the equation for equilibrium of the entire sutured repair (Eq. (4)).

The numerical solution of the state space ODEs was obtained using procedures built into the MATLAB (The Mathworks, Natick, MA) environment (ODE23s). Two sets of ODE functions with short and long term parameters were coded and the final displacement for the short term behavior was used as the initial condition for the long term ODE to evaluate the long term behavior.

## 3 Results

To investigate how the stiffness and anchor positions of rows of sutures can affect the force distribution over the repair, we varied the stiffnesses *k*_*n*_ and the lengths of the tendon segments secured between anchors. As described above, the relationship between the lengths of the tendon segments and the mechanical properties of these segments were chosen so that *L*_1_*/L*_2_ = *H*_1_*/H*_2_ = *K*_21_*/K*_11_ = *K*_22_*/K*_12_. To analyze the relationship between suture row and force distribution, suture stiffnesses were normalized with respect to *K*_*n*1_ + *K*_*n*2_ = 17.8 N/m, taken from the long term viscoelastic relaxation data.

When the force on the end of the tendon was ramped to a prescribed value, the forces in all three suture rows increased. Because the relaxation timescales for tendon (∼ 2 − 100 s) were slower than the 1 s muscle force ramp time, the distribution of forces over three rows of sutures varied over time (Fig. 4) in the manner that depended upon the stiffnesses of the sutures and their spacing.

**Figure 4:**
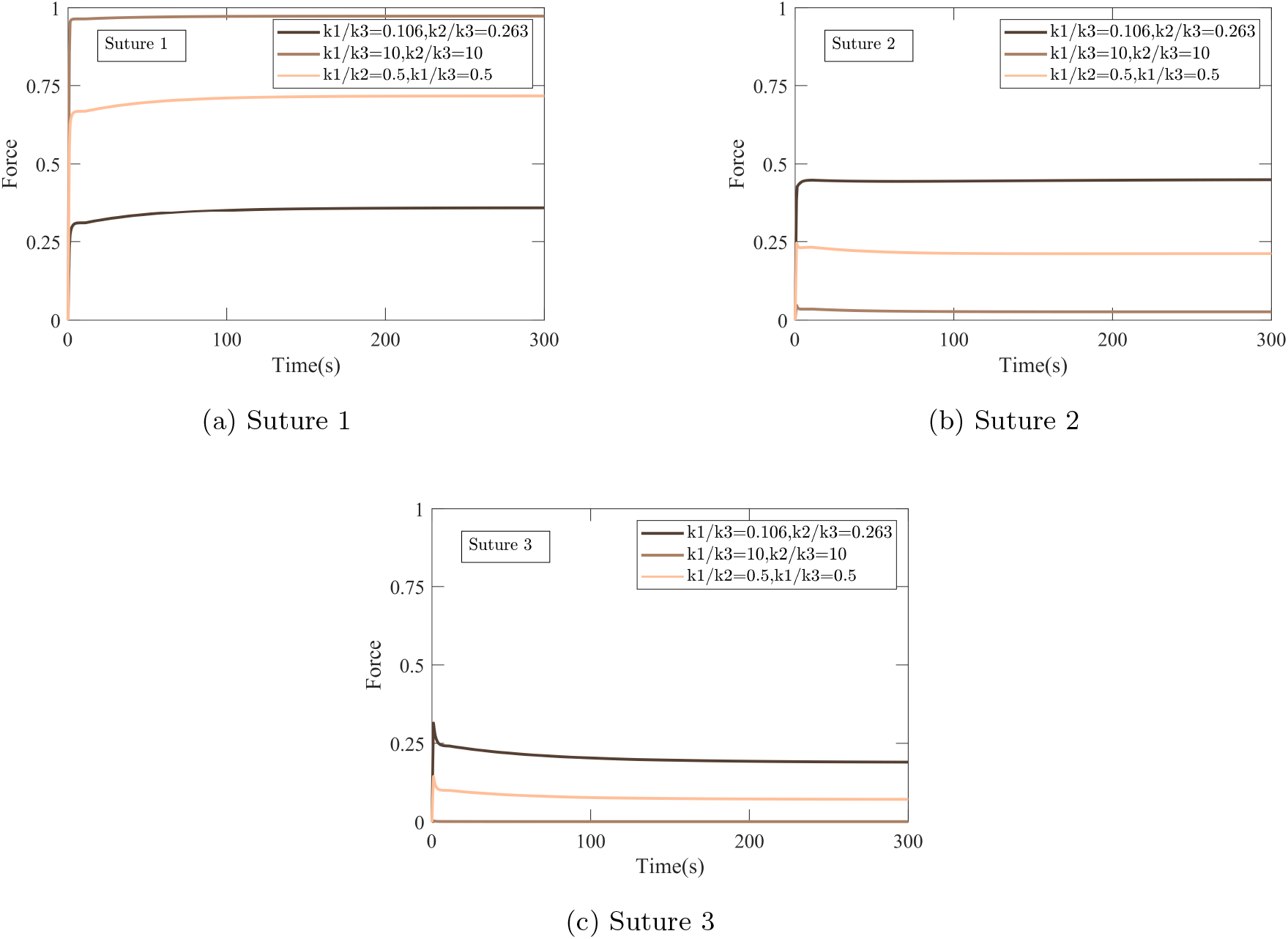
Forces on the sutures corresponding to three combinations of suture row stiffness ratio. (a) Suture row 1, (b) Suture row 2, (c) Suture row 3.

To evaluate the effects of suture row stiffness on the distributions of force, we plotted the suture row force distributions immediately after the application of the loading ramp (Fig. 5a) and the steady state distributions (Fig. 5b), which in all cases were reached within a few percent after 300 s of loading. In the case plotted in Fig. 5, suture rows were spaced equally, and tendon segments all had the same mechanical properties: *K*_11_ = *K*_21_ = *K*_31_, *K*_12_ = *K*_22_ = *K*_32_, and *H*_1_ = *H*_2_ = *H*_3_. The third suture row had fixed stiffness *k*_3_ = *K*_31_ + *K*_32_, the ratio *k*_1_*/k*_3_ = 0.106 was kept constant, and the stiffness *k*_2_ was varied. In cases where *k*_2_ and *k*_3_ were less stif than the stiffness of the tendon, suture row 2 was subject to the largest force. As the suture row stiffness increased, suture row 1 became subject to a higher force than suture row 2 at both the beginning of loading and in the steady phase.

**Figure 5:**
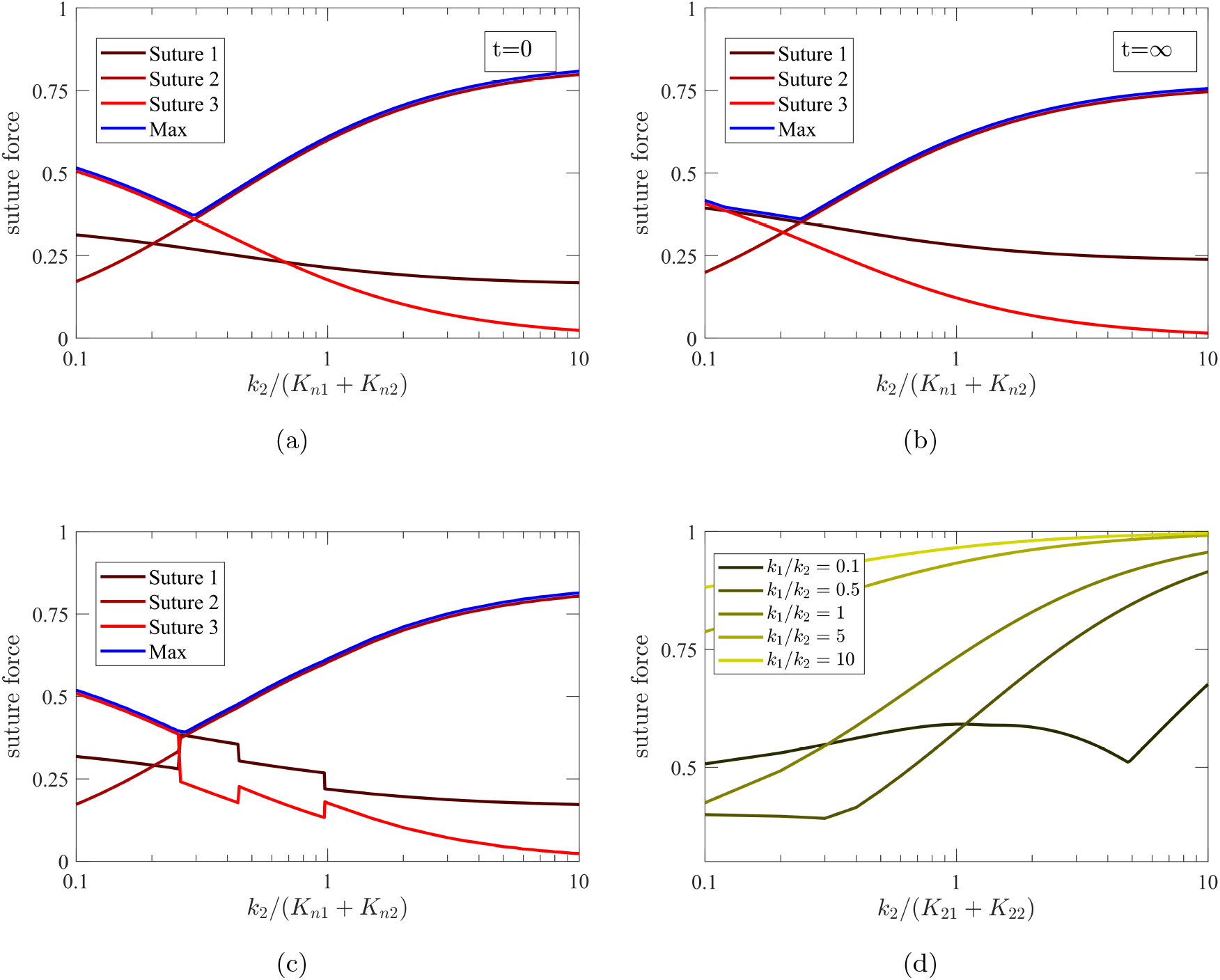
Suture row forces depended on time in ways that varied with the relative elastic stiffnesses of the suture rows. Here, *K*_11_ = *K*_21_ = *K*_31_, *K*_12_ = *K*_22_ = *K*_32_, and *H*_1_ = *H*_2_ = *H*_3_, while *k*_3_ = (*K*_31_ + *K*_32_) was kept constant, *k*_1_*/k*_3_ = 0.106, and *k*_2_ was varied. (a) Forces immediately after the 1 s loading ramp. (b) Forces at steady state. (c) Maximum forces in the suture rows over a loading cycle. (d) Maximum force among three suture rows in case where *k*_2_ = *k*_3_, with variable ratios of *k*_1_*/k*_2_, as a function of *k*_2_/(*K*_*n*1_ + *K*_*n*2_).

The maximum forces in each suture row over the entire loading duration did not vary monoton-ically with suture row stiffness due to transitions in the time at which peak loading was achieved (Fig. 5c). This plot was generated by finding the maximum force amongst all three suture rows for each set of stiffness parameters over the entire loading interval, then finding the forces in the other suture rows at that point in time. For lower values of *k*_2_, the peak force occurred immediately after loading in the third suture row, following the trend seen in Fig. 5a. For higher values of of *k*_2_, the peak occurred in the second suture row, following the trend observed in Fig. 5b. At the transition, the peak force occurred in the first suture row, as in Fig. 5b. A rapid transition between suture row 2 and suture row 3 could be seen, arising from the viscoelasticity of the tendon: immediately after the loading ramp, the force in suture row 2 was lower than that in suture row 3, but with time, relaxation in the tendon caused a shift in both the position and time of peak force as is observed in Fig. 5d. Results demonstrated that the stiffness ratio *k*_1_*/k*_2_ is critical to the force distribution. Variations of the values of *k*_1_*/k*_2_ can cause the maximum force in the suture rows to change by a factor of 2, without altering *k*_2_ and *k*_3_ values.

The specific set of parameters used for Fig. 5d was chosen for illustration. To evaluate the space of possible solutions more generally, a contour plot of peak force across the suture rows was plotted for varying values of *k*_1_*/k*_3_ and *k*_2_*/k*_3_, with *k*_3_ = *K*_31_ + *K*_32_ and suture row spacing fixed at *L*_1_*/L*_2_ = 0.72. In the contour plot (Fig. 6), a (dark blue) minimum force concentration could be seen for a specific combination of suture row stiffness. The space of solutions had a generally triangular appearance, with solutions with high *k*_1_*/k*_3_ corresponding to the peak force occurring in suture row 1, solutions with low *k*_1_*/k*_3_ and high *k*_2_*/k*_3_ corresponding to peak force occurring in suture row 2, and solutions with low *k*_1_*/k*_3_ and *k*_2_*/k*_3_ corresponding to peak force occurring in suture row 3. At the optimum, forces were distributed nearly equally across all three suture rows, with the maximum suture row force being 38.2% of the overall applied force.

**Figure 6:**
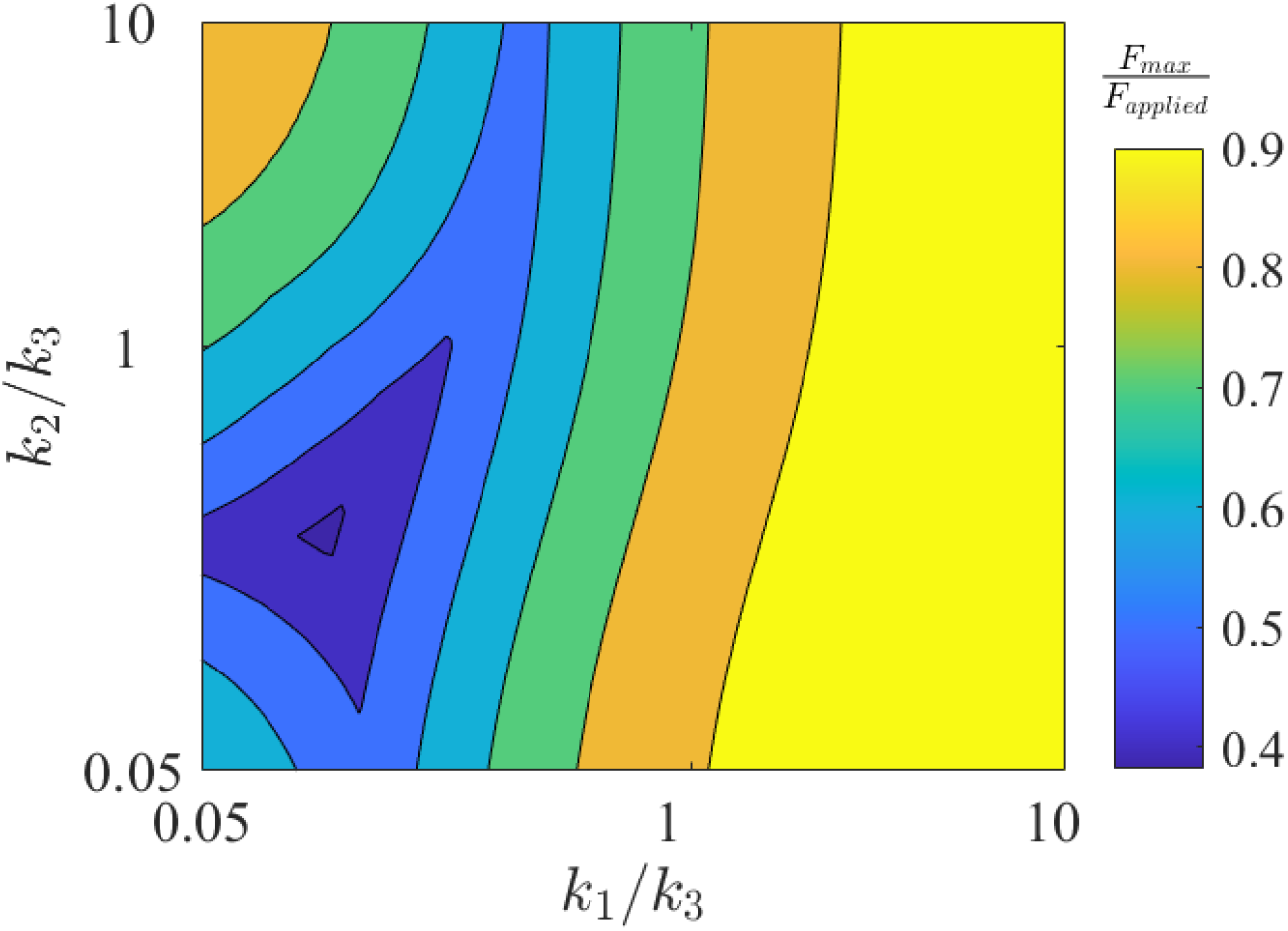
Effect of relative stiffness of tendon segments on the maximum force in the most loaded segment. Tendon segment length ratio *L*_1_*/L*_2_ at 0.72 and varying *k*_1_*/k*_2_, *k*_1_*/k*_3_. The vertical colorbar to the right of the color map shows the range of the maximum force on suture rows with forces varying from high (yellow color) to low (blue color).

To demonstrate the effect of suture row spacing on force balancing, for one set of parameters, the analysis was performed using the optimum suture row stiffness ratio observed in Fig. 6 (*k*_1_*/k*_3_ = 0.106 and *k*_2_*/k*_3_ = 0.263). As distance *L*_1_ between the first two suture rows increased relative that of the distance *L*_2_ between the second two suture rows (increasing *L*_1_*/L*_2_), the fraction of the total forced carried by suture row 1 increased, while the fractions carried by suture rows 2 and 3 both decreased (Fig. 7). When the first two suture rows were close (low *L*_1_*/L*_2_ ratio), the peak load moved to the second suture row, and the minimum moved to first suture row. For an optimal length ratio near *L*_1_*/L*_2_ =0.72, the forces the suture rows were nearly balanced.

**Figure 7:**
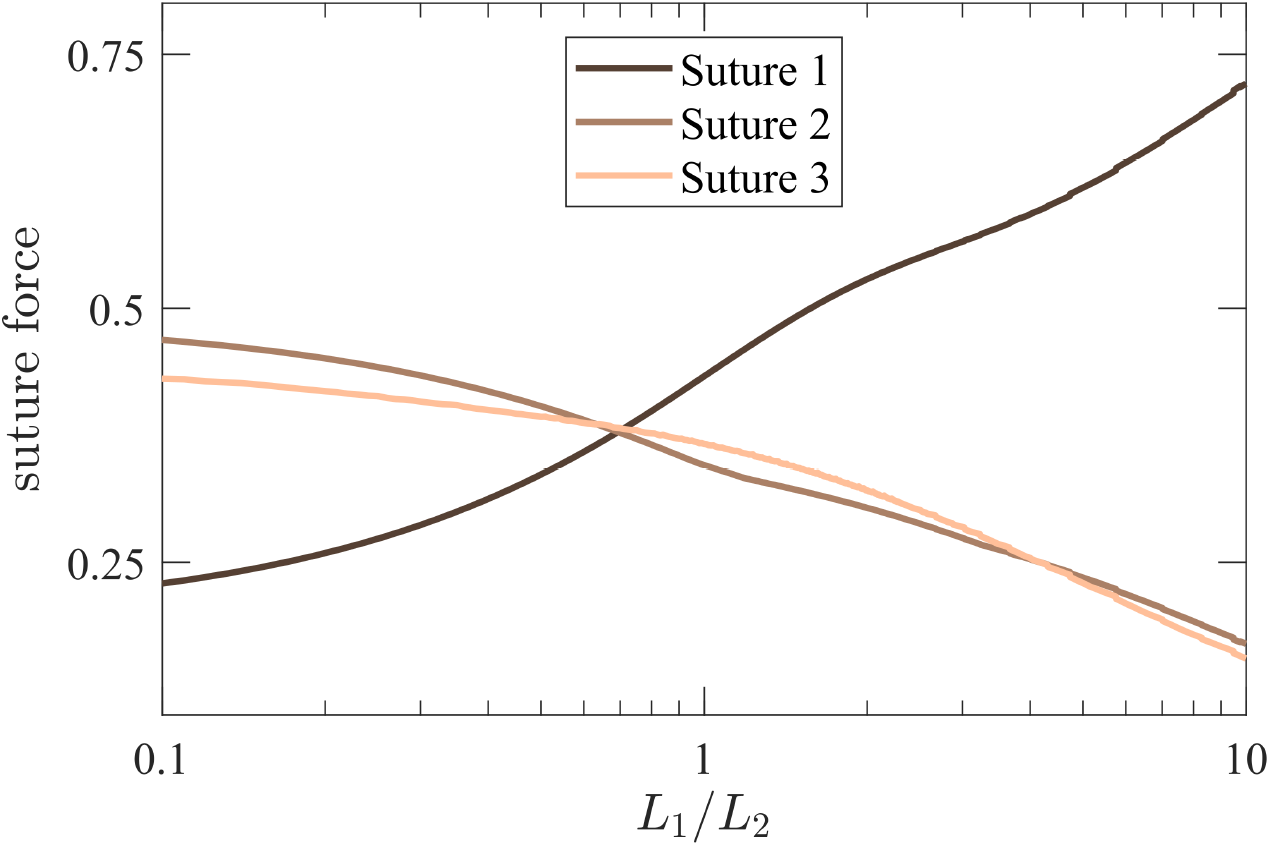
Suture row locations versus the suture row forces in parameter studies *k*_1_*/k*_3_ = 0.106 and *k*_2_*/k*_3_ = 0.263

To illustrate the inter-related effects of suture row spacing and suture row stiffnesses on force balancing, *L*_1_*/L*_2_ and *k*_2_*/k*_3_ were varied, while *k*_3_ = *K*_31_ + *K*_32_ and the stiffness ratio *k*_1_*/k*_3_ = 0.106 were constant (Fig. 8). The resulting contour plot showed a (dark blue) minimum of 38.2% as the peak fraction of the total force taken by a suture row. Once more, the space of solutions had a generally triangular shape, solutions with high *k*_2_*/k*_3_ and low *L*_1_*/L*_2_ corresponding to high peak force on suture row 2, solutions with low *L*_1_*/L*_2_ and *k*_2_*/k*_3_ corresponding to peak force occurring in suture row 3, and solutions with high *L*_1_*/L*_2_ corresponding to peak force occurring in suture row 1. A discontinuity in slope was evident in the contours in the upper left of the plot, indicative of a shift of force balance over time associated with viscoelastic effects.

**Figure 8:**
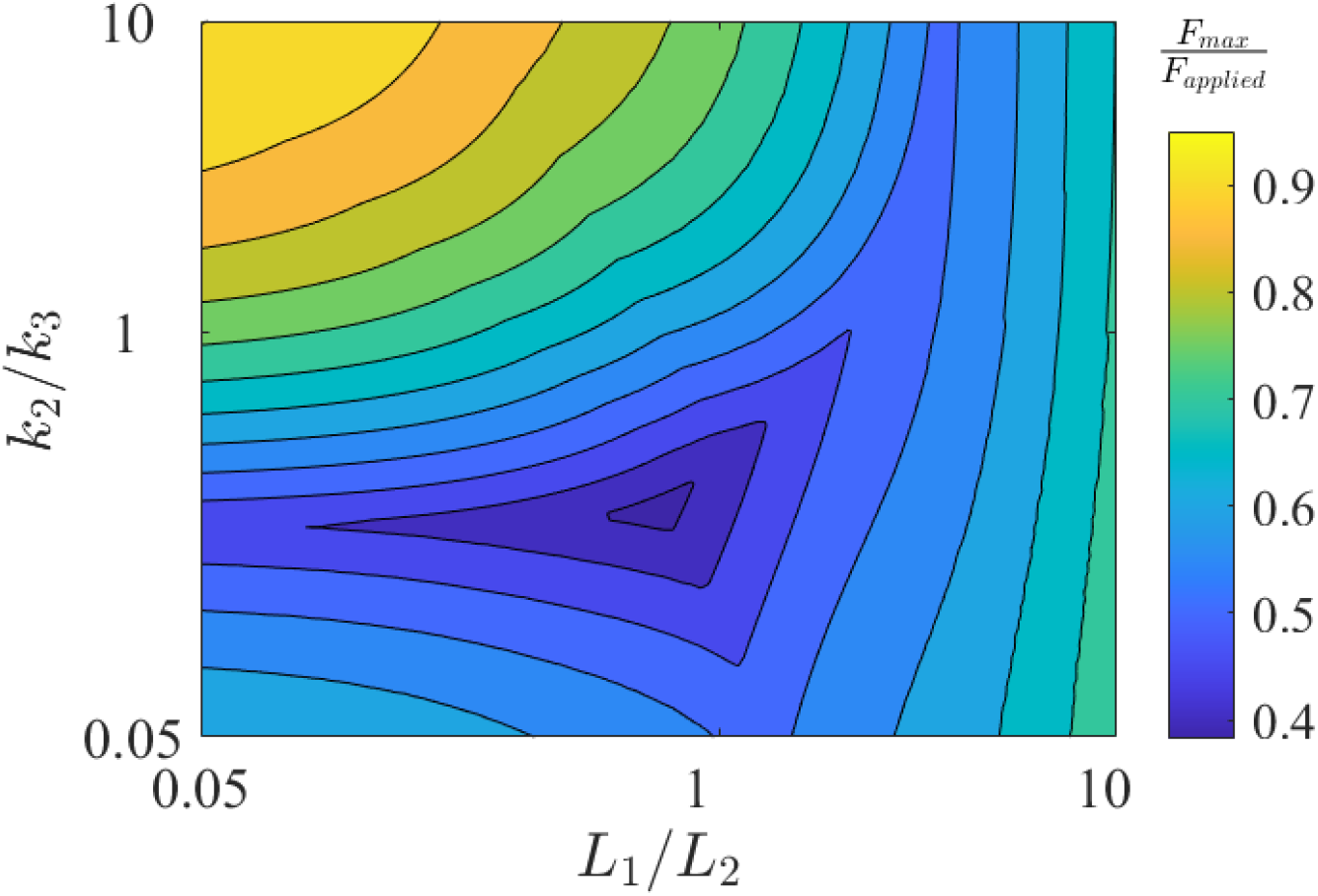
Peak fraction of the applied force acting on a row of sutures as a function of suture row stiffness and tendon segment length ratio, with *k*_1_*/k*_3_ = 0.106.

## 4 Discussion

Biomaterials, including tendons and ligaments, are in general viscoelastic. In the present study, we considered the distribution of forces in suture rows and tendon in a tendon tear repair employing three suture rows, and found that tissue viscoelasticity led to a rebalancing of forces over time. The initial balance of forces and its variation over time was a strong function of the relative stiffnesses and anchor spacing of the suture rows. Results suggest that, by adjusting these parameters, a surgeon may be able to increase repair strength balancing forces over the three rows of sutures. While the strength of typical sutures is not affected by such optimization, the issue is important to the strength of the repair because suture pull-out forces are related to the stress concentration at the anchoring points of the sutures, and balancing forces thus reduces the peak stress within a tendon.

Using the solution obtained by the viscoelastic model introduced in the paper, the difference between short-term and long-term force duration responses was observed in Figs. 5a and 5b. In equally spaced suture rows of equal stiffness, the largest force is transmitted through suture row 1 (the outermost suture row), while the other suture rows are underloaded. A benefit of the optimization of the repair was clearly evident from Fig. 4 where variations of the suture row stiffnesses lead to a much more balanced force distribution compared to less efficient designs. Varying the stiffness of suture rows 1 (outermost suture row) and 2 (middle suture row) (Fig. 7) balanced forces to the point that the maximum force in any suture row over time was 38.2% of the applied external force, representing a significant reduction compared to alternative “non-optimum” designs.

The model contains many idealizations that need to be mentioned. Suture rows were modeled as linear elastic springs, while the tendon sections between the anchoring points were analyzed using a three-parameter linear viscoelastic model, accounting for their elastic and viscous properties. The response of the repair idealized as explained here was considered, concentrating on the relative stiffness and positioning of three suture rows to achieve a desirable force distribution. Even better results may be possible by combining sutures with an adhesive layer bonding tendon to bone ([11, 24]). Similar ideas of combined bolted-bonded attachments have been considered in repairs of aerospace structures (e.g., [25]) and proven effective.

## 5 Conclusions

Conventional rotator cuff repair techniques introduce high stress concentrations in the tendons that often leads to repair failure. Some of these stress concentrations can be alleviated by altering the anchor suture spacing and the stiffness of the suture rows. In the present study, we considered an alternative design employing suture rows with different stiffnesses and unequal anchor suture spacing to achieve a more even distribution of the forces. Moreover, we accounted for viscoelastic effects in the tendon. The optimization conducted for a three-suture row model suggested that relatively simple adjustments to current repair techniques can result in a much more favorable distribution of forces across the suture rows, reducing stress concentrations in the tendon.

